# Fixation probabilities in network structured meta-populations

**DOI:** 10.1101/2021.07.21.453198

**Authors:** Sedigheh Yagoobi, Arne Traulsen

## Abstract

The effect of population structure on evolutionary dynamics is a long-lasting research topic in evolutionary ecology and population genetics. Evolutionary graph theory is a popular approach to this problem, where individuals are located on the nodes of a network and can replace each other via the links. We study the effect of complex network structure on the fixation probability, but instead of networks of individuals, we model a network of sub-populations with a probability of migration between them. We ask how the structure of such a meta-population and the rate of migration affect the fixation probability. Many of the known results for networks of individuals carry over to meta-populations, in particular for regular networks or low symmetric migration probabilities. However, when patch sizes differ we find interesting deviations between structured meta-populations and networks of individuals. For example, a two patch structure with unequal population size suppresses selection for low migration probabilities.

## Introduction

Evolutionary graph theory^1^ analyses the evolutionary dynamics in network-structured populations, where individuals are located on the nodes of a graph and can replace each other via the links. This approach can be motivated by several biological systems, from the spread of cancerous mutations through colonic crypts to the invasion of ecosystems structured by rivers. One of the major goals of evolutionary graph theory^1^ is to assess the effect of underlying population structure on the fixation probability (the probability of ultimate fixation of a mutant)^2–6^ and the fixation time^7–10^. An important aim is to find an optimized structure to speed up or slow down the spread of a newly arising mutant^11–13^ .

However, to apply such models, a change in perspective is often necessary: In many biological applications, the nodes correspond to small populations and not to single individuals^14^. Replacement of individuals via the links is then exchanged with migration where the immigrant displaces one from the resident individuals. This leads to network-structured meta-populations, where a network is formed by individual populations connected via migration. An important question that arises in the application of evolutionary graph theory is thus whether the results derived for networks of individuals carry over to networks of small populations.

Traditionally, such meta-populations have been analyzed extensively in ecology, where they correspond to fragmented habitats^15–17^. The dynamics of the meta-population is driven by the exchange of individuals between subpopulations. Nevertheless, the focus of these studies is usually quite different, as they aim to address complex questions arising in ecology by asking for the impact of such population structure. Also in population genetics, the fixation probability in subdivided populations (meta-populations) has been investigated extensively, see e.g. the study by Maruyama in 1970^18^, who has shown that the fixation probability is not altered by the subdivision of a population into partially isolated patches under certain assumptions. Later, other types of structured populations and alternative modes of selection and evolutionary dynamics have been discussed in this context^19–23^. However, the focus in the context of population genetics is not on the network structure of the meta-population structure. Typically, in these systems individuals from every patch have the freedom to migrate to all the other patches, whereas in our system individuals migrate to a small subset of adjacent patches.

Also, evolutionary game dynamics and the evolution of cooperation have been studied in subdivided populations^2, 24–30^. Yet, with few exceptions, see e.g.^31^, these studies assume a well-mixed population of patches.

Here, we extend the typical models of evolutionary graph theory to network-structured meta-populations. In contrast to previous work that heads into this direction, our focus is on very basic models of populations of fixed size and a small number of patches. We discuss limiting cases and show how they can be addressed with the available analytical tools. We show that the isothermal theorem holds for any meta-populations with the regular structure. Since an exact solution for the fixation probability is not feasible when the network structure is complicated, an approximation for small migration probability is used to compute the fixation probability. Finally, we discuss the two-patch meta-population and meta-star, where several leaf patches are connected only to one central patch, when the migration probability is high.

## Model

We consider a finite population of size *N* distributed in a network of *M* patches (Fig. 1). The local population size of patch *j* is constant and given by *N_j_*. Each individual can reproduce either within its own patch or place its offspring into an adjacent patch.

**Figure 1.**
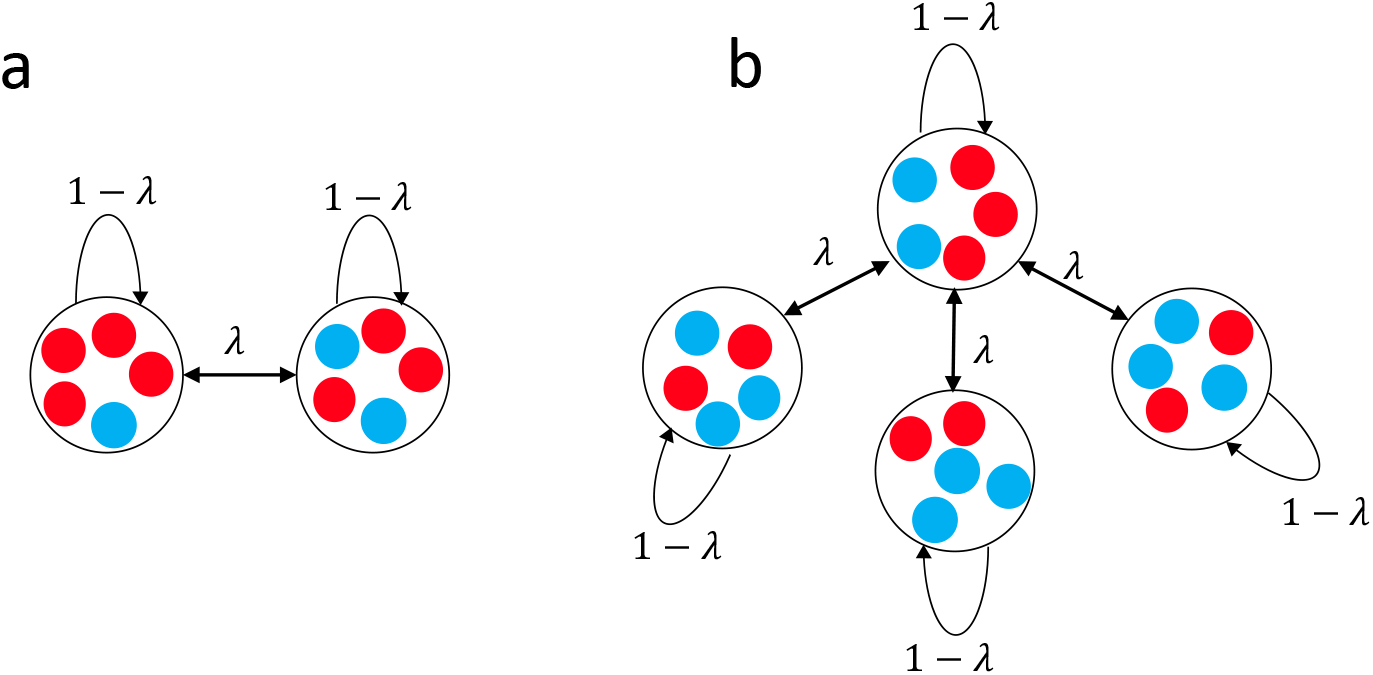
Network of populations; the population consists of well-mixed subpopulations coupled through migration (a) two-patch meta-population with local patch size *N*_1_ = *N*_2_ = 5, (b) meta-star with *M* = 4 patches and local population size *N*/*M* = 5.

We consider two types of individuals, mutant and wild-type. We start with a wild-type population of size 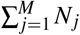. In the first step, a random wild-type individual turns into a mutant. We work with a standard Birth-death process with global selection for birth and random local death^1, 32^: In each time step, one individual is selected from the whole population at random, but proportional to its fitness, to produce identical offspring. Afterwards, the newborn replaces one of the individuals within its patch, including its mother, with probability 1 – *λ*, or it replaces one of the individuals in a random adjacent patch with probability *λ*. To be more precise, suppose that patch *i* is connected to a set of other patches *V* . If *α* is an individual in patch *i* and *β* is an individual in patch *j* ≠ *i*, then the probability that an offspring of *α* replaces *β* is equal to 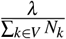, where *N_k_* is the population size in patch *k*. If *α* and *β* are individuals in the same patch, *i*, the same probability is 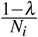.

Our goal is to examine how likely this new mutant takes over the whole population and how this probability changes with the migration probability and the network structure.

For our model, the comparison with the fixation probability within a single patch of a well-mixed population is important. Therefore, we first recall how to calculate this quantity. Suppose the population of size *N* consists of wild-types with fitness 1 only. If a new beneficial or deleterious mutant with fitness *r* appears in the population, the evolution of the population will lead to the fixation or extinction of the mutant. As we adopt a Birth-death process the fixation probability, 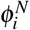, starting with *i* mutants is determined by^33–35^

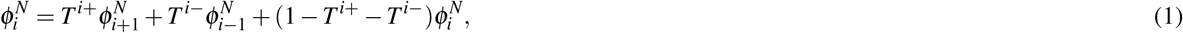

where 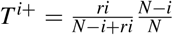 is the probability that the number of mutants increases from *i* to *i* + 1 and 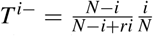 is the probability that the number of mutants decreases from *i* to *i* – 1. With this recursive relation and the boundary conditions 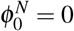 and 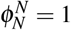, the fixation probability of a single mutant equals

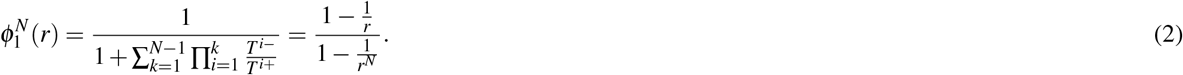

Note that for *r* → 1, the fixation probability converges, 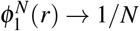. This classical result will be an important reference point for our further considerations, we will denote it by 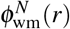 in the remainder (where “wm” stands for well-mixed).

## Results

### Regular structures and isothermal theorem

For networks where each node represents a single individual, the isothermal theorem of evolutionary graph theory shows that the fixation probability is the same as the fixation probability of a well-mixed population if the temperature distribution is homogeneous across the whole population^1^. The temperature of a node defined as the sum over all the weights leads to that node. This theorem extends to structured meta-populations for any migration probability *λ*: If the underlying structure of the meta-population that connects the patches is a regular network and the local population size is identical in each patch, the temperature of all individuals is identical, regardless of the value of the migration probability. Therefore, the fixation probability in a population with such a structure is the same as the fixation probability in a well-mixed population of the same total population size 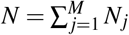, given by 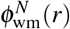.

### Small migration regime

If the migration probability is small enough such that the time between two subsequent migration events 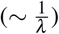 is much longer than the absorption time within any patch, then at the time of each migration event we may suppose that the meta-population is in a homogeneous configuration^20, 26^. In other words, the low migration regime is an approximation in which we neglect the probability that the meta-population is not in a homogeneous configuration at the time of migration events. We define a homogeneous configuration of the meta-population as a configuration in which in all patches either all individuals are mutants, or all are wild-types.

Therefore, instead of having 2*^N^* states, where *N* is the population size, the system has only 2*^M^* states, where *M* is the number of patches. Thus, we can calculate the fixation probability exactly as in the case of a standard evolutionary graph model where each node represents a single individual. Let us use *T^j→k^* for the probability to replace the individual in node *k* by an offspring arising in node *j* in a network of individuals. To move towards a network of patches, we only need to substitute *T^j→k^* with the product of the migration probability of a mutant from patch *j* to patch *k* and the fixation probability of the mutant, 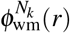,

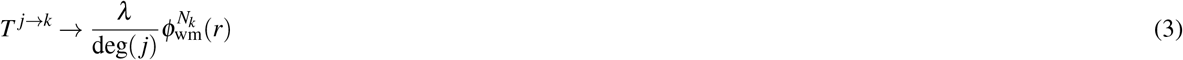

where deg(*j*) is the degree of node *j* to take into account that the mutant can move to different patches. For the case of a wild type individual migrating into a mutant patch, we need to replace *T^j→k^* instead by

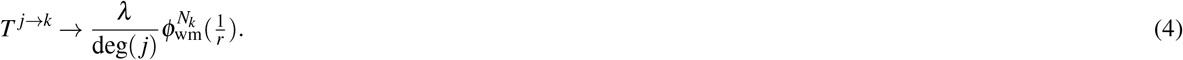

Since *λ* appears in both probabilities of increasing and decreasing the number of mutants, it drops out of the resulting equation for the fixation probability.

#### Two-patch meta-population

The simplest non-trivial case is the fixation probability in a two-patch meta-population with different local size for small migration probability *λ*. If the migration probability *λ* is very small, a new mutant first needs to take over its own patch and only then the first migrant arrives in the second patch. To be more precise, the time between two migration events has to be much higher than the typical time that it takes for the migrant to take over the patch or go extinct again^36^. In this case, we can divide the dynamics into two phases: A first phase in which a mutant invades one patch and a second phase in which a homogeneous patch of mutants invades the whole meta-population. Assume a new mutation arises in patch 1. Only if this mutant reaches fixation in patch 1, it also has a chance to reach fixation in patch 2. When patch 1 consists of only mutants and patch 2 consists of only wild-types, there are two possibilities for the ultimate fate of the mutant:

i. Eventually, the offspring of one mutant selected from patch 1 for reproduction will migrate to patch 2 and reach fixation there. The wild-type goes extinct. This happens with probability 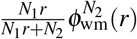.
ii. Eventually, the offspring of one wild-type selected from patch 2 for reproduction will migrate to patch 1 and the mutant goes extinct. This occurs with probability 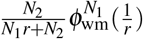.

Therefore, the probability that a single mutant arising in patch 1 reaches fixation in the entire population is

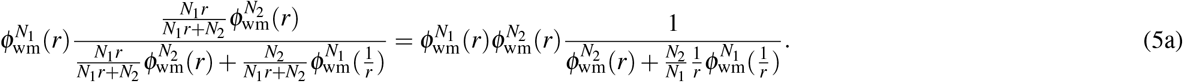

Similarly the probability that a mutant arising in patch 2 takes over the whole population equals

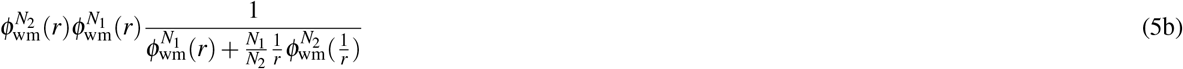

If we assume that the mutant arises in a patch with a probability proportional to the patch size, the average fixation probability 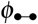 in a two patch population for small migration probability is the weighted sum of Eqs. 5a and 5b,

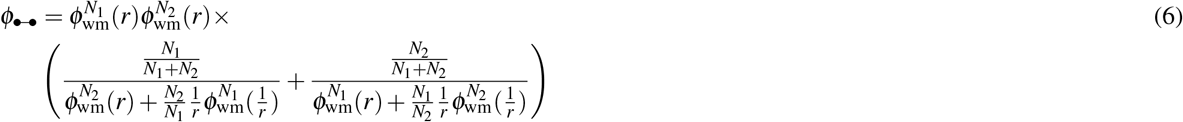

In the case of neutrality, *r* = 1, we recover 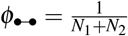 – the fixation probability in a population of the total size of the two patches. For identical patch sizes, *N*_1_ = *N*_2_, Eq. (6) simplifies to

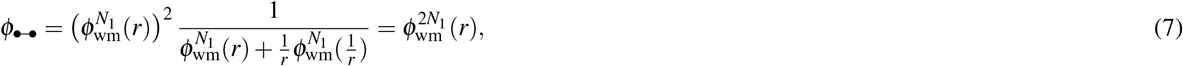

where the simplification to the fixation probability within a single population of size 2*N*_1_ reflects the validity of the isothermal theorem.

For *N*_1_ ≠ *N*_2_, we approximate Eq. (6) for weak and strong selection. Let us first consider highly advantageous mutants, *r* >> 1. In this case, we have 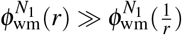 and thus we can neglect the possibility that a wild-type takes over a mutant patch if patch sizes are sufficiently large. The probability 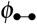 then becomes a weighted average reflecting patch sizes. For identical patch size *N*_1_ = *N*_2_ = *N*/2, it reduces to 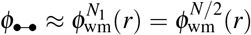. In other words, taking over the first patch is sufficient to make fixation in the entire population certain. For patches of very different size, *N*_1_ >> *N*_2_, we have *N* ≈ *N*_1_ and find 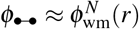, which implies that fixation is driven by the fixation process in the larger patch, regardless of where the mutant arises. Note that there is a difference between the case of identical patch size and very different patch size . The case of highly disadvantageous mutants, *r* << 1, can be handled in a very similar way.

Next, we consider weak selection, *r* ≈ 1. We can approximate the fixation probability as 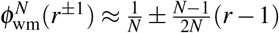. with this, we find

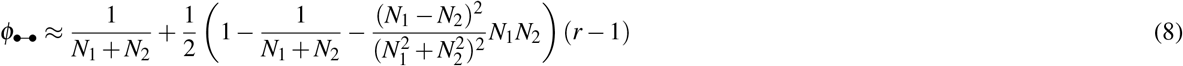

For identical patch size *N*_1_ = *N*_2_ = *N*/2, this reduces to

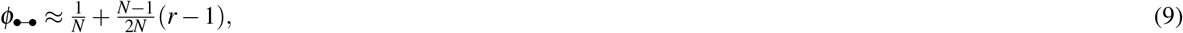

which is the known result for a single population of size *N* = *N*_1_ + *N*_2_. When patches have very different size, *N*_1_ ≫ *N*_2_ such that *N* ≈ *N*_1_, we recover the same result. Thus, the difference between the fixation probability of a two-patch meta-population with identical patch size and the fixation probability of a two-patch meta-population with very different patch size that we found for highly advantageous mutants is no longer observed for weak selection.

When migration probabilities become larger, our approximation is no longer valid and we need to rely on numerical approaches. Fig. 2 illustrates the difference between the fixation probability of a two-patch structure meta-population and the equivalent well-mixed population of size *N*_1_ + *N*_2_ when migration is low using Eq. 6 and comparing with the numerical approach in^37^.

**Figure 2.**
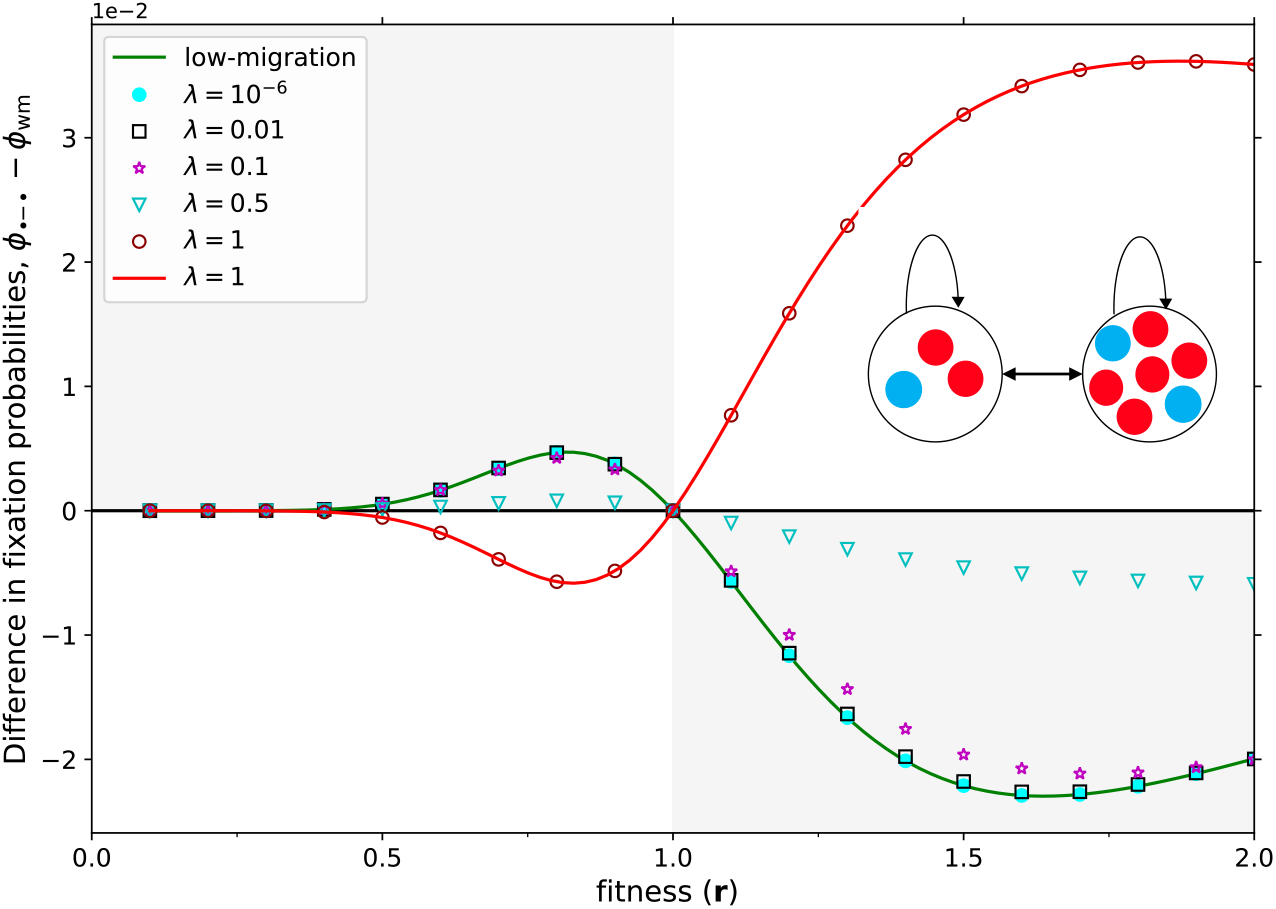
The difference between the fixation probability of a two-patch meta-population and a well-mixed population of the same size. The two patch sizes are *N*_1_ = 3 and *N*_2_ = 7 as a function of fitness. Lines show analytical results for low migration probabilities (Eq. (6)) and migration probability *λ* = 1 (Eq. (21)). Symbols show numerical results based on a transition matrix approach^37^. The numerical result and analytical result for low migration probability and high migration match perfectly. In the low migration regime the two-patch meta-population is a suppressor of selection, indicated by the fact that the symbols are never in the area with a white background. However, in the high migration regime (*λ* = 1), where simulations and analytical results again match, the two-patch meta-population is an amplifier of selection. The fixation probability for *λ* = 1 is obtained analytically using the Martingale approach discussed by Monk^39^.

While the fixation probability of the two-patch meta-population is very close to the fixation probability of the well-mixed population^38^, a close inspection reveals an interesting property: For low migration probabilities and *N*_1_ ≠ *N*_2_, the two patch structure is a suppressor of selection in the original sense of Lieberman et al.^1^: For advantageous mutations, *r* > 1, it decreases the fixation probability, whereas for disadvantageous mutations, *r* < 1, it increases the fixation probability compared to the well mixed case. For weak selection, we show this analytically: For *r* ≈ 1, we can write

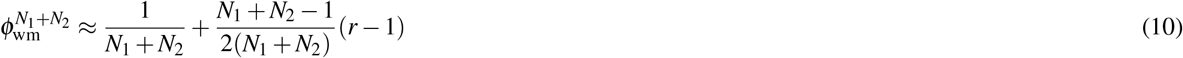

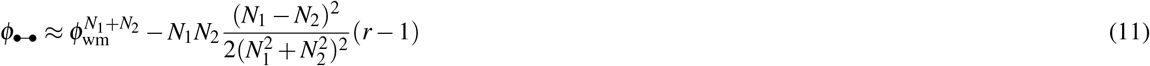

While the difference to the well mixed case vanishes for *N*_1_ = *N*_2_ in first order in *r* – 1, the fixation probability of the two patch structure is larger for *r* < 1 and smaller for *r* > 1. Thus, under weak selection the two patch structure with *N*_1_ ≠ *N*_2_ is a suppressor of selection.

While the structure remains a suppressor of selection for most values of the migration probability *λ* , Fig. 2 reveals that for very large *λ* it becomes an amplifier of selection.

#### The meta-star

Here, we approximate the fixation probability *ϕ*_⋆_ of a meta-star with *M* – 1 leaves for low migration probability. For simplicity, we assume that all patches are of the same size 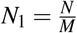 and omit the notation for patch size. As long as the migration probability is sufficiently low, such that before the next migration the immigrant gets fixed or lost, patches tend to be homogeneous. We denote the number of homogeneous mutant patches among the leaves by *j* and use a lower index to represent the state of the central patch, which is either occupied by wild-types (∘) or by mutants (•). The number of homogeneous mutant patches increases in two ways:

i. the center is occupied by mutants and is selected for birth and its offspring migrates to one of the peripheral homogeneous wild-type patches and reaches fixation in that patch,

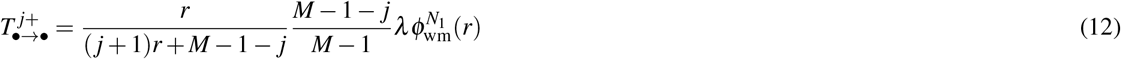

where 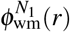 determines the fixation probability of a mutant in a local population,
ii. the center is occupied by wild-types and one of the homogeneous mutant leaves is selected for birth and its offspring migrates to the center and gets fixed there,

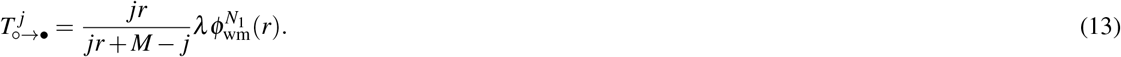

Note that the number of homogeneous mutant leave nodes cannot increase if the center is occupied by wild-type individuals, i.e. 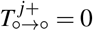. Similarly, the number of homogeneous mutant leave nodes cannot decrease if the center is occupied by mutants, i.e. 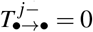. Thus, the number of homogeneous mutant patches can decrease in two ways,

i. the center is occupied by wild-types and is selected for birth and its offspring migrates to one of the homogeneous mutant leaves and gets fixed there,

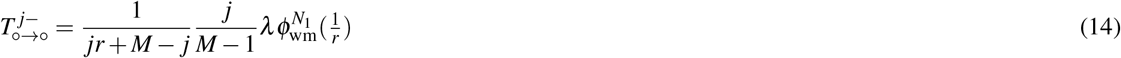

where 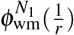 is the fixation probability of a wild-type in a local mutant population, or
ii. the center is occupied by mutants and one of the leaves is selected for birth and its offspring migrates to the center and gets fixed there,

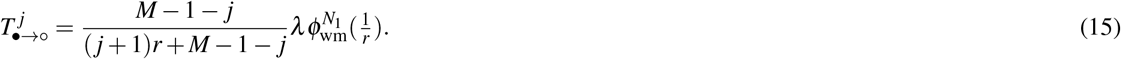

For the star graph, depending on where the mutant emerges, two different fixation probabilities are defined: The fixation probability when a single mutant emerges in the center 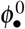 and the fixation probability when a single mutant emerges in one the leaves, 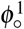. Following the same arguments as in^40^, we find

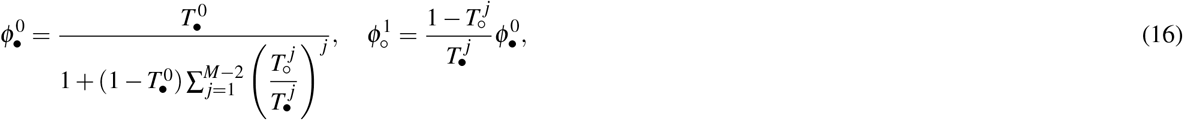

where the probability to leave the state with a wild-type patch in the center and *j* mutant patch leaves is

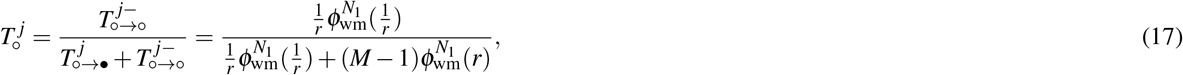

and the probability to leave the state with a mutant patch in the center and *j* mutant patch leaves is

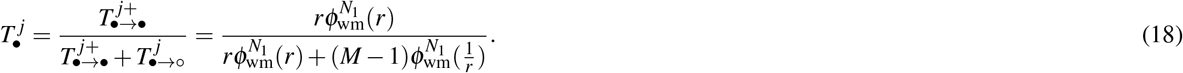

Note that the two probabilities 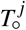 and 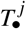 are independent of *j* in our particular case. Thus, also their ratio 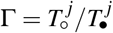 is independent of *j*, which makes our calculation of 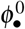 easier.

Evaluating Eqs. (16), we find the average fixation probability in the entire patch structured meta-population starting from a single homogeneous mutant patch,

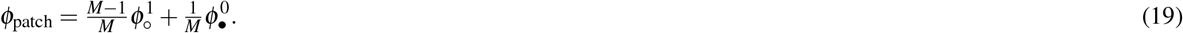

Therefore, the fixation probability of the whole population equals

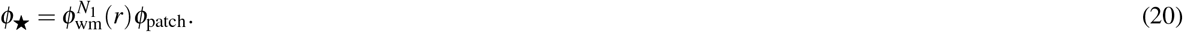

Fig. 3 illustrates the fixation probability of a meta-star as a function of fitness. Numerical solutions for low migration agree very well with the low migration approximation. According to this plot, the meta-star is an amplifier of selection in the low migration regime – similar to the star network of individuals^1^. A numerical investigation of Eq. (20) reveals that this result carries over to larger *N*_1_ as well. For any value of *M* and *N*_1_ between 1 and 100, we find that the star network of patches amplifies selection. However, as expected from earlier work^38^, the extent of amplification becomes smaller with growing population size.

**Figure 3.**
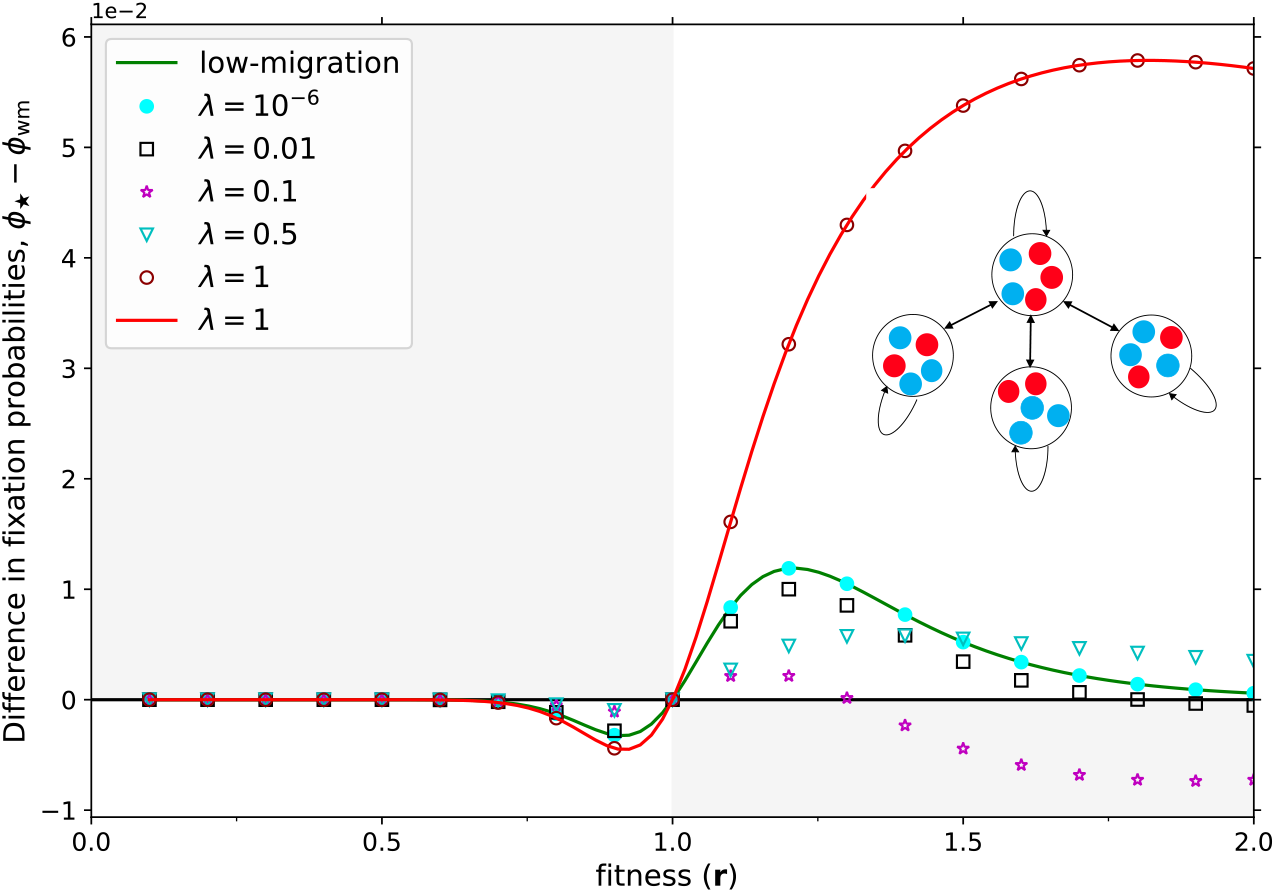
The difference between the fixation probability of a meta-star and the equivalent well-mixed population of size with *M* = 4 patches and identical local size in each patch *N*_1_ = *N/M* = 5 for different fitness values. Lines show analytical results for low migration probabilities (Eq. (20)) and migration probability *λ* = 1 (Eq. (21)) symbols show numerical results based on a transition matrix approach^37^ for *λ* = 10^−6^, *λ* = 0.01, *λ* = 0.1, and *λ* = 0.5. For migration probability *λ* = 10^−6^, we observe an almost perfect agreement, the low migration result serves as a good approximation. The fixation probability in *λ* = 1 is obtained analytically using the Martingale approach. For extremely high and low migration probability the meta-star acts as an amplifier of selection (such that the lines only pass through the white shaded area of the plot) while in the intermediate migration regime shows a very different behavior where it could be an amplifier or a suppressor of selection depending on the fitness value.

Using the same approach as we used for the two patch meta-population, we find that the meta-star is an amplifier for small and high migration probability, but not in between. For intermediate migration probability, it is only a piecewise amplifier^41, 42^ and does not fall into one of the originally defined categories, see Fig. 3.

The meta-star in low migration is equivalent to the “star of islands” discussed by Allen et al.^43^. In their study for death-Birth updating they found that the comparison of the size of the hub to the size of the leaves makes a determinative difference. When the leaves are larger, the structure amplifies under weak selection; when the hub is larger, it suppresses under weak selection. When the hub and leaves are the same size, the structure acts as a “reducer”, meaning that it lessens the fixation probability for all r not equal to 1 (termed “suppressor of fixation” elsewhere^32^). Doing the same comparison in the whole range of selection, we find that when leaves and hub are of the same size, the meta-star is an amplifier under Birth-death updating. When the size of the leaves exceeds the size of hub it is a transient amplifier and finally, when the size of the hub exceeds size of the leaves it amplifies selection.

### High migration probability

For moderate migration probabilities, it is challenging to calculate the fixation probability. However, in the case of the maximum possible migration probability, *λ* = 1, the two-patch meta-population and meta-star transform to complete bipartite graphs: In the two-patch meta-population, every offspring will be immediately moved to the other patch. In the meta-star, the offspring of individuals in the center node will be placed in a random leaf, whereas the offspring of the individuals in the leaf nodes will be placed in the center. Thus, the meta-star can be thought of as a bipartite graph in which one part is made out of all leaf nodes and the other part out of the center.

The fixation probability of complete bipartite graphs has been calculated analytically previously^39, 42, 44^ using the specific features of Martingales. The probability to reach fixation if the initial mutant arises either in patch 1, 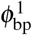 or patch 2, 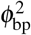 are

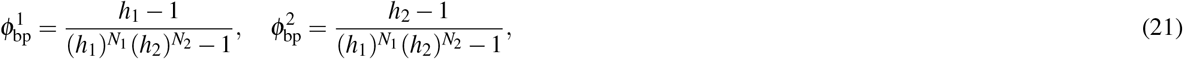

where 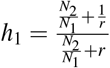 and 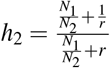. The average fixation probability is

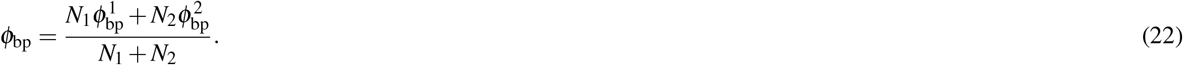

In the case where one of the patch size is much larger than the other, *N*_2_ ≫ *N*_1_, the fixation probability converges to the fixation probability of a star graph 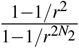.

As discussed above, a star-structured meta-population in *λ* = 1 can be reduced to a complete bipartite graph. As a result the fixation probability of a star meta-population with *M −* 1 leaves and population size *N* such that the population distributing homogeneously in all the patches is obtained by replacing *N*_1_ with 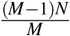 and *N*_2_ with 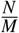 in Eq. 21 and 22.

As shown in Fig. 2 and Fig. 3 both the two-patch meta-population and meta-star are amplifiers of selection for *λ* = 1. It has been proven in^45^ that a complete bipartite amplifies selection for weak selection. This can also be seen from a Taylor expansion of the difference between Eq. (22) and the corresponding result for the well mixed population at *r* = 1, which leads to

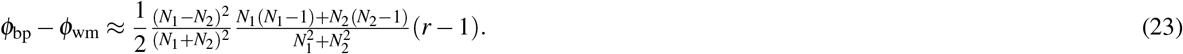

This quantity is positive for *r* > 1 and negative for *r* < 1, such that the structure is an amplifier of selection for weak selection. Eq. (22) reveals this fact holds for the whole range of selection strength. If we have a fixed population of size *N* on a complete bipartite graph, for fitness values *r* > 1 the minimum fixation probability occurs when the two patch sizes are identical, *N*_1_ = *N*_2_ = *N*/2. Similarly, for fitness values *r* < 1 the maximum fixation probability occurs when the two patch sizes are identical, *N*_1_ = *N*_2_ = *N*/2.

Since a complete bipartite graph with identical patch size is an isothermal graph and its fixation probability is the same as the fixation probability of a well-mixed population, we conclude that any complete bipartite graph is the amplifier of selection when *N*_1_ ≠ *N*_2_. This result is implicitly contained in^39, 44^, but deserves special attention: It implies that a graph can be turned into the amplifier if we enforce a very large degree of exchange of individuals between patches. Combined with the observation that many graphs of individuals are amplifiers of selection^32^, it suggests that it may be easier to construct amplifiers of selection than suppressors of selection in undirected networks^5, 46^.

## Discussion

Here, we have extended evolutionary graph theory from graphs of individuals to evolutionary graph theory of populations. We have investigated how the structure and migration probability influence the probability to fixation. For regular networks and patches of identical size, any evolutionary graph of populations has the same fixation probability as the well mixed population for any migration probability – a result that follows directly from the isothermal theorem^1^.

However, for non-regular networks or patches of different size, this is no longer the case and the dynamics depends on the migration probability. If the migration probability is small enough, such that the time to fixation in one patch is small compared to the time between two subsequent migration events, we can use time-scale separation to approximate the fixation probability. Using this approximation, we show that the two-patch meta-population suppresses selection whenever the two patches have different size. Based on the same approximation, we have shown that the meta-star amplifies selection. In the high migration regime, the two-patch meta-population and the meta-star can be viewed as complete bipartite graphs. Evolutionary dynamics in bipartite graphs has been studied in^39, 44^. Both the two-patch meta-population and the meta-star are amplifiers of selection in this high migration regime.

Here, in order to be as close to the original ideas of evolutionary graph theory and the popular Birth-death updating as possible, we have focussed on meta-populations in which selection is global. Thus, individuals compete across the entire meta-population for reproduction. Another possible condition would be when the competition is local i.e. among the individuals belonging to the same patch. This would correspond to a process where a site becomes available and there is local competition to fill it, similar to death-Birth dynamics in networks of individuals^43, 47^. The choice of this dynamics can have massive consequences on the amplification or suppression properties of a graph^32^. In general, structured meta-populations allow many additional selection processes that are still to be analyzed.

## Methods

Depending on the network and the range of migration we adopt three different approaches to calculate the fixation probability:

i. In the very low migration probability we implement an analytical approximation for the fixation probability using the time scale separation between migration and fixation in a single patch.
ii. For intermediate migration probability we compute the fixation probability numerically using the transition matrix based on approach published in^37^. The entries of the transition matrix represent the transition probability between different possible states. The number of states depend upon the symmetry of the network.
iii. For very high migration probability we calculate the fixation probability analytically using the Martingales introduced in^44, 48^.

We model the evolutionary dynamics using the Moran process. The code to reproduce our figures is available at https://github.com/s-yagoobi/fixation-probability.

## Acknowledgements

We thank Stefano Giaimo and Nikhil Sharma and Florence Bansept for constructive discussions on this subject.

## Author contributions statement

A.T. designed the study. S. Y. designed and implemented the code and performed the numerical analysis. S.Y. and A. T., analyzed the simulation data and developed analytical calculations. Both authors wrote the manuscript together.

